# Genome-wide selection signatures reveal widespread synergistic effects of culture conditions and temperature stress in *Drosophila melanogaster*

**DOI:** 10.1101/2021.11.09.467935

**Authors:** Burny Claire, Nolte Viola, Dolezal Marlies, Schlötterer Christian

## Abstract

Experimental evolution combined with whole-genome sequencing is a powerful approach to study the adaptive architecture of selected traits, in particular when replicated experimental populations evolving in opposite selective conditions (e.g. hot vs. cold temperature) are compared. Nevertheless, such comparisons could be affected by environmental effects shared between selective regimes (e.g. laboratory adaptation), which complicate the interpretation of selection signatures. Here, we used an experimental design, which takes advantage of the simplicity of selection signatures from founder populations with reduced variation, to study the fitness consequences of the laboratory environment (culture conditions) at two temperature regimes. After 20 generations of adaptation at 18°C and 29°C, strong genome-wide selection signatures were observed. About one third of the selection signatures can be either attributed to temperature effects, laboratory adaptation or the joint effects of both. The fitness consequences reflecting the combined effects of temperature and laboratory adaptation were more extreme in the hot environment for 83% of the affected genomic regions, fitting the pattern of larger expression differences between founders at 29°C. We propose that evolve and resequence (E&R) with reduced genetic variation allows to study genome-wide fitness consequences driven by the interaction of multiple environmental factors.

## INTRODUCTION

Ecological genetics aims to characterize the interaction of organisms with their environment. Of particular interest is the characterization of adaptive responses, which are specific to a given habitat. Many approaches have been pursued to study the genetic basis of local adaptation (Savolainen et al., 2013, Tiffin & Ross-Ibarra, 2014, Whitlock, 2015, Hoban et al., 2016, Lorant et al., 2020). Allele frequency differences between populations from different environments are particularly powerful to test for correlation between genetic variation and environmental variables (Coop et al., 2010, Günther & Coop, 2013), and are widely applied to the analysis of clinal variation (Rako et al., 2007, Kolaczkowski et al., 2011, Fabian et al., 2012, Bergland et al., 2016, Calfee et al., 2020). Despite being conceptually appealing, this approach faces several challenges. Historical demographic events, such as bottlenecks or admixture, may generate confounding signals, possibly resulting in false positives/negatives (Thornton & Jensen, 2007, Pavlidis et al., 2010, Lohmueller, 2014, Lotterhos & Whitlock, 2015, Johri et al, 2020). Furthermore, estimating covariance between allele frequencies and environment is difficult as i) identifying and/or measuring the relevant environmental variables is challenging since many abiotic factors are correlated ( Mittler, 2006, MacColl, 2011) and ii) selection can vary over time (e. g. Bergland et al., 2014, Behrman et al., 2018, Grainger et al., 2021).

Experimental evolution, in particular laboratory natural selection, allows to study adaptive responses in a controlled laboratory environment (Burke & Rose, 2009, Garland & Rose, 2009, Kawecki et al., 2012, Schlötterer et al., 2015). Exposing a mixture of genotypes to a monitored stressor, the adaptive response can be measured through time in replicate populations, combined with next-generation sequencing (Evolve and Resequence (E&R); Turner et al., 2011, Schlötterer et al., 2014, Long et al., 2015). While many experimental evolution studies rely on truncating selection to determine the genotypes contributing to the next generation (Turner et al., 2011, Turner & Miller, 2012, Griffin et al., 2017, Hardy et al., 2018, Gerritsma et al., 2019), laboratory natural selection builds on fitness differences between genotypes upon exposure (Garland & Rose, 2009) and hence provides a closer fit to adaptation and competition processes occurring in the wild (Hsu et al., 2021).

A major challenge for the interpretation of molecular selection signatures comes from the few recombination events during the laboratory experiment resulting in strong linkage disequilibrium (Nuzhdin & Turner, 2013, Tobler et al., 2014, Franssen et al., 2015). Strong linkage reduces the efficiency of natural selection as a consequence of Hill-Robertson effect (Hill & Robertson, 1966, Roze & Barton, 2006). Starting with many different founder genotypes (Baldwin-Brown et al., 2014, Kofler & Schlötterer, 2014, Kessner & Novembre, 2015, Vlachos & Kofler, 2019) and using the selected haplotype blocks as the selective unit rather than individual SNPs (Franssen et al., 2017, Barghi et al., 2019, Otte & Schlötterer, 2021) may partially overcome the lack of resolution. Nevertheless, increasing the number of founders will increase the pool of adaptive variants and consequently the number of beneficial genotypic combinations to reach the trait optimum (Yeaman, 2015, Barghi et al., 2019, Barghi & Schlötterer, 2020, Laruson et al., 2020). One proposed solution to study the selective response of highly polygenic traits builds on reducing the genetic variation in the founder population (Sachdeva & Barton, 2018, Burny et al., 2021, Langmüller et al., 2021). Assuming that even the use of only two haplotypes provides sufficient segregating variation to adapt to rapid thermal change, we focused on laboratory adaptation as an environmental factor common to two different temperature regimes. We used 18°C, a putatively non-stressful temperature regime since the two founder genotypes of our experiment showed very similar gene expression profiles at 18°C (Chen et al., 2015, Jaksic & Schlötterer, 2016). In contrast, 29°C is a very stressful temperature regime, close to the maximal temperature at which *D. melanogaster* populations can be maintained (Hoffmann, 2010). We observed a very strong selection response across the entire genome. About one third of the genomic regions responded either only to temperature, laboratory conditions, or exhibited a significant joint effect of both stressors. Our results demonstrate the importance of the combined effects of different environmental factors.

## MATERIALS AND METHODS

### Experimental set-up

We used the Oregon-R and Samarkand strains inbred by Chen et al. (2015), and maintained since then at room temperature. The three replicates of both experimental evolution cages were set up in parallel, each with a census size of 1,500 flies and accidentally with a starting frequency of 0.3 for the Oregon-R genotype (0.7 for the Samarkand genotype) - rather than 0.5, as described in Burny et al, 2021. Briefly, all replicates were then maintained for 20 generations at either constant 29°C or constant 18°C in dark conditions before sequencing. 300 adults were transferred every generation to one of five bottles for two days of egg laying. After egg laying, all adults were removed and frozen. The egg lay resulted in a high density of larvae. Hence, we transferred a mixture of larvae and food to two fresh food bottles. Adults collected 8-32 hours after eclosure of the first flies from all bottles were mixed to avoid population substructure. 300 adults from each vial started the next generation.

### DNA extraction, library preparation, sequencing

Whole-genome sequence data for the parental Oregon-R and Samarkand strains are available in Burny et al, 2021. The evolved replicates in generation F20 were sequenced using Pool-Seq: genomic DNA was prepared after pooling and homogenizing all available individuals of a given replicate in extraction buffer, followed by a standard high-salt extraction protocol (Miller et al., 1988). For the samples in the 29°C experiment, barcoded libraries with a targeted insert size of 480 bp were prepared using the NEBNext Ultra II DNA Library Prep Kit (E7645L, New England Biolabs, Ipswich, MA) and sequenced on a HiSeq 2500 using a 2×125 bp paired-end protocol. For the samples in the 18°C experiment, we used the same library preparation protocol, but with a target insert size of 280 bp, and 2×150 bp reads were sequenced on the HiSeq X Ten platform.

### Allele frequency tracking

We previously established a catalogue of parental SNPs (Burny et al., 2021). Briefly, a parental SNP was defined as a (nearly) fixed difference between parental lines with a 0/0 (1/1) genotype in the Samarkand parent and 1/1 (0/0) genotype in the Oregon-R parent at the marker position, conditioning for a frequency of the alternate allele lower than 0.05 (if 0/0) or higher than 0.95 (if 1/1) for a final list of 465,070 SNPs; 401,252 and 63,818 SNPs on the autosomes and the X chromosome, respectively, equivalent to 1 SNP every 271 bp on the autosomes and 363 bp on X. The same processing and mapping steps were applied at 29°C and 18°C described in (Burny et al., 2021). The allele frequency have been obtained after converting processed BAM files from pileup (*samtools mpileup -BQ0 - d10000*; version 1.10; (Li et al., 2009)) to sync files (using PoPoolation2 *mpileup2sync.jar*; (Kofler et al., 2011)). We then tracked the allele frequency at F20 of the Oregon-R allele in 3 replicates at 29°C (replicates 1,2,3 in Burny et al, 2021) and 3 replicates at 18°C. The subsequent analyses have been performed with R (version 4.0.4; (R Core Team 2020)) and most panels have been generated with the ggplot2 R package (Wickham, 2016). We retained SNPs measured at both temperatures, leading to a total of 100,283, 89,929, 107,119, 103,760, 72, 63,766 SNPs on 2L, 2R, 3L, 3R, 4 and X. Because the average coverage at the marker SNPs differs between both temperatures (12, 11, 9× at 18°C and 123, 107, 133× at 29°C), we down-sampled the 29°C coverage values to 12× by drawing the coverage at each locus from a Poisson distribution of mean 12 and then applying binomial sampling with a sample size set to the sampled coverage to mimic Pool-Seq sampling noise (Taus et al, 2017). In order to both limit noise in allele frequency measurements and to take linkage into account, the allele frequency values are averaged in non-overlapping windows of size *w*=50, 250 or 500 SNPs for a total of 8,021, 1,603, 801 measurements on the autosomes (2 and 3) and 1,275, 255, 127 on X for each window size respectively, where the last window of each chromosome, containing fewer than *w* SNPs. Windows of size *w*=50, 250 or 500 SNPs correspond to 13.6 [12.8; 14.4], 67.8 [59.7; 76.0] and 135.6 [115.8; 155.4]kb on average for the autosomes and 18.2 [16.6; 19.7], 90.5 [81.4; 99.5] and 180.9 [162.1; 199.8]kb for X. The 95% confidence intervals have been obtained by the mean +/−1.96 SE, with SE standard error. The main results are represented at 250-bp level. A window position *i* is defined by its center ((right-left)/2). By convention, if the Oregon-R allele frequency at F20 is higher (lower) than its initial frequency of 30% (70%), the Oregon-R (Samarkand) allele increased in frequency and the allele frequency change (AFC) is positive (negative).

### Comparing the response between the 18°C and 29°C selection regimes

We classified the AFC of each window after 20 generations as non-significantly deviating from neutrality or presenting a selection signal. In order to test deviation from neutrality, we performed 100 neutral simulation runs using MimicrEE2 (Vlachos & Kofler, 2018). The neutral simulations mimic the experimental set-up, *i.e*. starting with 30% of Oregon-R flies over 1,500 flies, using three replicates and the same marker SNPs providing the *D. melanogaster* recombination map (Comeron et al., 2012) updated to version 6 of the reference genome using the Flybase online Converter (https://flybase.org/convert/coordinates; accessed in July 2020). For each simulation run, we computed the average AFC over the three replicates per window. Per temperature and per chromosome, an empirical p-value per window *w* (p_w_^18°C neutral^ or p_w_^29°C neutral^) is calculated as the fraction of AFC values higher (lower) than the empirical AFC when the observed AFC is positive (negative) divided by the total number of average AFC values. We finally applied a Benjamini-Hochberg correction per chromosome (p.adj_w_^18°C neutral^ and p.adj_w_^29°C neutral^). If a window presents a selection signal, it either favors the same parental allele at both temperatures (with a change in magnitude or not) or different alleles - for example the Oregon-R allele at 29°C (AFC_w_^29°C^>0) and the Samarkand allele at 18°C (AFC_w_^18°C^<0). To check which scenario is more likely, we fitted a simple linear model (LM) for each window *w*, with AFC as response and temperature as fixed categorical explanatory factor, where *α_w_^intercept^* corresponds to 18°C-reference level and *α_w_^temperature^* is the contrast between 29°C and 18°C. We extracted the corresponding p-value (p_w_^LM^) and applied a Benjamini-Hochberg correction per chromosome on the non-neutral windows (p.adj_w_^LM^). A significant window is classified as displaying a change in magnitude with the temperature favoring the same parental allele (*α_w_^intercept^* and *α_w_^temperature^* of same sign) or a different allele (*α_w_^intercept^* and *α_w_^temperature^* of different sign). For a given False Discovery Rate (FDR) threshold, a genomic window w is then classified in one of the following 6 classes: “drift only”, “change 18°C only”, “change 29°C only”, “no temperature effect”, “different magnitude” and “different direction” (see Table SI 1 for logical conditions on windows affectation to each class). We then recorded the fraction of windows affected in each of the 6 classes for different values of FDR (5%, 10%, 15%) per chromosome and averaged genome-wide (GW). We also computed the autocorrelation per chromosome and per replicate using the *acf* R function; the autocorrelation at a given step *k* is defined as the correlation between windows at positions *i* and *i+k*, where *k* is called the lag. We eventually recorded the distance where a significant decrease in autocorrelation at a 5% threshold (below 1.96/√*n*, *n* the number of windows), *i.e*. a rough proxy of linkage equilibrium, is reached.

### Ancestral gene expression re-analysis

We used ancestral gene expression values at 18°C and 29°C for each genotype (Chen et al., 2015). The parental gene expression is reported as the log2-transformed fold change of expression of the Samarkand genotype relative to the Oregon-R genotype expression used as a reference, noted logFC S/O. In order to correlate parental gene expression and allele frequency changes, we computed the AFC per gene as the average of AFC of parental markers located within the gene. To that aim, we needed to convert the genes position to the updated version of the *D. melanogaster* GTF annotation (v6.36). We downloaded the gene conversion IDs from FlyBase using “wget ftp://ftp.flybase.net/releases/current/precomputed_files/genes/fbgn_annotation_ID_*.tsv.gz” the 25^th^ November 2020. Over 7,853 gene expression values, remained 7,844 genes for which the conversion was possible. We then computed per gene the average AFC of all SNPs within the entire genic region (exons, introns and UTRs) over a total of 7,751/7,844 genes due to the sparse distribution of marker SNPs with on average 36 markers (median of 12) per gene. We first searched for the presence of any genome-wide correlations between the logFC S/O differential (logFC S/O 29°C - logFC S/O 18°C) and the AFC differential (AFC 29°C - AFC 18°C) paired by gene, measured by the Spearman correlation coefficient ρ. Assuming that correlation, if it exists, might be caused by a subset of genes, we also computed ρ coefficients for an increased number of top genes (by subsets of 50 genes) either ranked by the absolute logFC S/O differential or by the absolute AFC differential. To assess if the obtained trend, an exponential decrease of ρ with an increasing number of genes was more often seen than under a random ordering of the genes, we computed for each set of top x genes and for each ranking, the 95^th^ quantile of 100 randomly chosen set of x genes. Eventually we performed a transcription factor binding sites (TFBS) enrichment analysis until 5kbp up-stream of each gene for the top 50 genes either ordered by decreasing logFC S/O at 18°C (48 genes present in the motifs database) or by logFC S/O at 29°C (44 genes present in the motifs database) using the RcisTarget bioconductor package (version 1.6.0; (Aibar et al., 2017)). The motifs database was downloaded from https://resources.aertslab.org/cistarget/databases/drosophila_melanogaster/dm6/flybase_r6.02/mc8nr/gene_based/dm6-5kb-upstream-full-tx-11species.mc8nr.feather the 20^th^ November 2020. Enrichment was defined using the default enrichment score of 3 and the number of motifs associated to a TFs was reported for each analysis.

## RESULTS

We exposed two genotypes, Samarkand and Oregon-R, to two different environmental stressors, laboratory adaptation and temperature. Two E&R experiments shared the same laboratory environment, but differed in temperature regime. Three replicate populations were maintained for 20 generations at either 18°C or 29°C. Genome-wide allele frequencies of genotype-specific marker SNPs were determined by Pool-Seq (Schlötterer et al., 2014). Because genotype-specific alleles start at the same frequency in all replicates and only few recombination events were expected during the experiment, we averaged the allele frequencies in non-overlapping windows of 250 consecutive SNPs to obtain reliable allele frequency estimates. This strategy is supported by the high autocorrelation of neighboring SNPs, up to a distance of 6.7Mb (Fig SI 1). We inferred selection by contrasting the allele frequencies of the Oregon-R genotype at the start of the experiment (30%) to those after 20 generations, relative to simulated frequency changes under neutrality. A positive allele frequency change (AFC) indicates that the Oregon-R allele increased in frequency.

After 20 generations marked allele frequency changes occurred at both temperature regimes (Fig 1A). The three replicate populations of each temperature regime showed a strikingly parallel selection response as indicated by the shaded area corresponding to +/− one standard deviation around the mean of the 3 replicates (Fig 2A). Overall, Oregon-R alleles were more likely to increase in frequency than Samarkand alleles, with 90% and 80% of the windows displaying positive AFC at 18°C and 29°C respectively.

**Figure 1.**
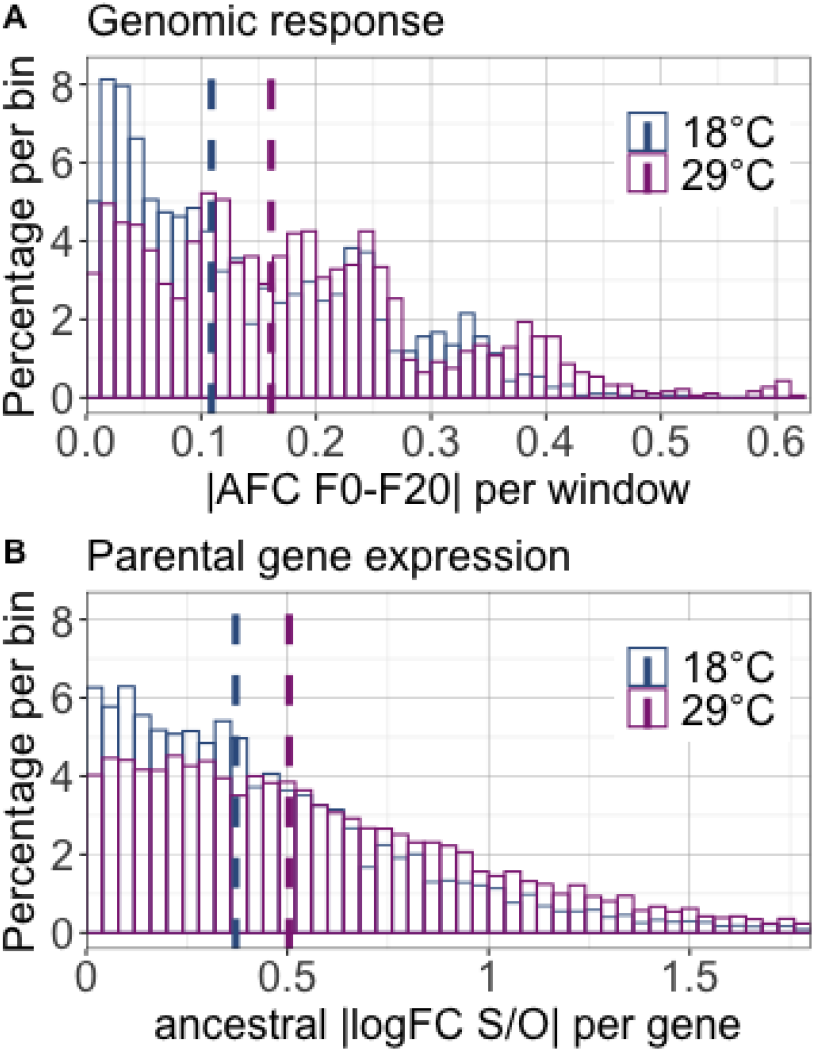
Differences between the parental genotypes at 18°C (blue) and 29°C (purple). Histograms of absolute allele frequency change of the Oregon-R allele between F0 and F20 (|AFC F0-F20|) for non-overlapping windows of 250 SNPs (A) and the absolute log2-fold difference of expression between Samarkand and Oregon-R genotypes (|logFC S/O|) per gene (B). The percentage of windows (A) and genes (B) in each of the 50 equally-spaced bins (bin size: 0.0125 (A), 0.04 (B)) is reported on the y-axis. The dashed lines represent the median absolute allele frequency change at 18°C and 29°C (0.11 at 18°C and 0.16 at 29°C, A; paired Wilcoxon one-sided test p-value=1.3×10^−40^) and median absolute logFC S/O (0.37 at 18°C and 0.50 at 29°C, B; paired Wilcoxon one-sided test p-value=1.8×10^−152^). For the sake of clarity, the x-axis of panel B is bounded at 1.8, which corresponds to 1.5 times the maximum inter-quartile range of the gene expression data. The full histogram is shown in Fig SI 2.

**Figure 2.**
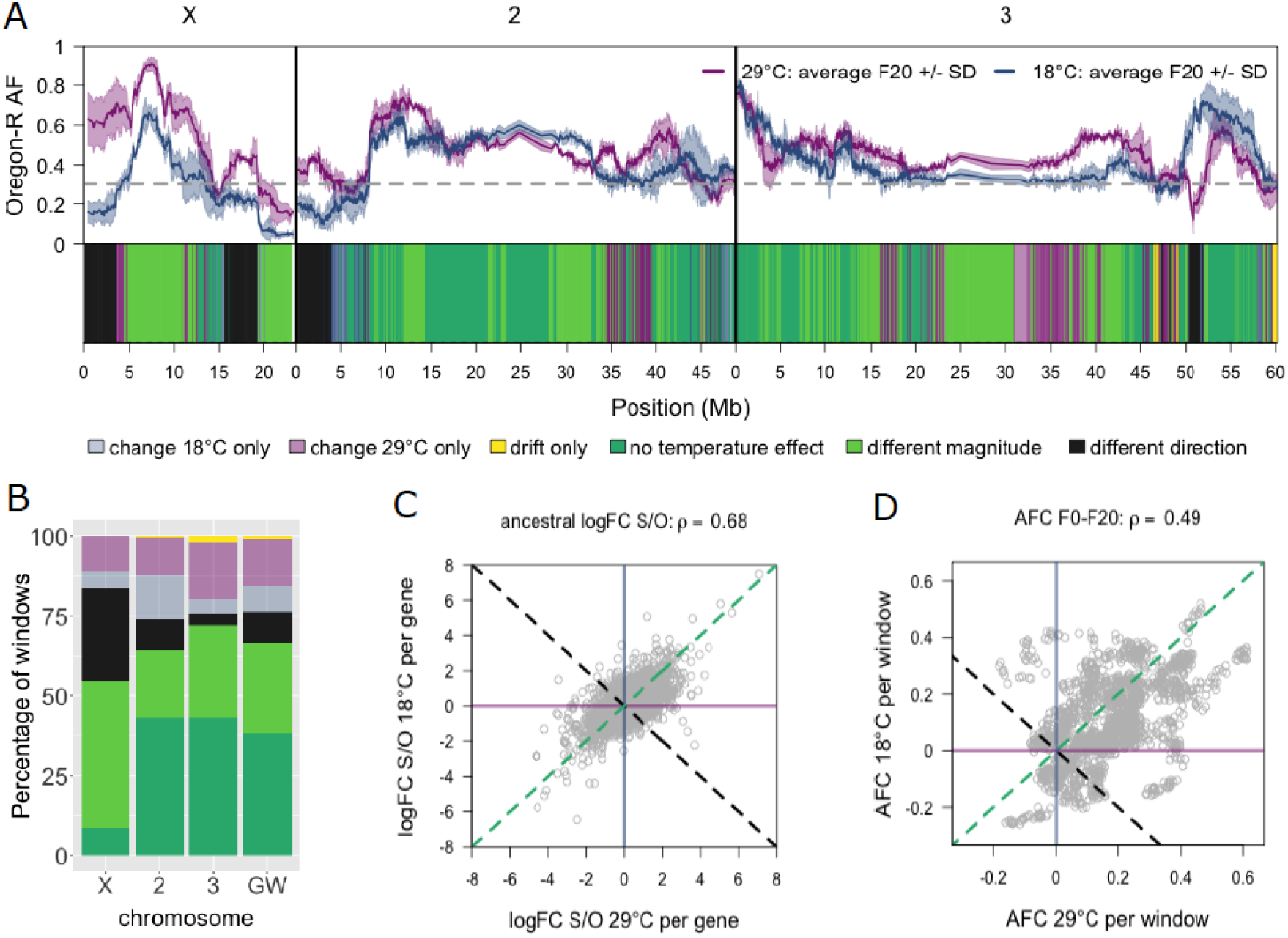
A) *Top*. Genome-wide allele frequencies after evolving for 20 generations at two temperature regimes. The frequency of the Oregon-R allele is averaged for non-overlapping windows of 250 SNPs (solid line) +/− one standard deviation (shaded area) at 18°C (blue) and 29°C (purple). *Bottom*. Each window is classified (see Methods) in one of the 6 color-coded classes depending of the AFC between 18°C and 29°C: change at 18°C only (light blue), change at 29°C only (light purple), drift only (yellow), no temperature effect (dark green), different AFC magnitude but same direction of effect (light green), opposite alleles increase at 18°C and 29°C (black). B) Percentage of the genomic windows per class (defined for an FDR threshold of 10% per chromosome and averaged genome-wide (GW)). Scatterplots of the logFC S/O (C) and AFC (D) at 18°C (y-axis) and 29°C (x-axis). We reported the Spearman ρ correlation coefficients.

Given that the *Drosophila* populations were adapting to two different environmental stressors, laboratory environment and temperature, it is possible to evaluate their individual and joint effect on the selection response across the entire genome. We characterized the selection response by classifying windows changing more in frequency than expected under neutrality in each of the temperature regimes (Fig 2A,2B, Table SI 1). On the one hand, the direction of the selection response, *i.e*. the increase in frequency of the Oregon-R or Samarkand alleles, differed for 10% of the windows between the two temperatures (Fig 2, black). 8% (Fig 2, light blue) and 14% (Fig 2, purple) of the windows displayed a significant allele frequency change relative to drift, only at either 18°C or 29°C respectively. On the other hand, a similar allele frequency change was observed for 38% (Fig 2, dark green) of the windows in the two experiments, which we attribute to laboratory adaptation only. In total, roughly 2/3 of the genome responded only to one of the two environmental stressors. Nevertheless, a remarkable large fraction of windows showed a significant combined effect of the two environmental stressors. 28% (Fig 2, light green) changed in the same direction, but at a different magnitude between temperatures. This pattern of frequency change indicates that temperature modulates the adaptive response to the selective force, common to both experiments. For most of these windows (83%), the most extreme allele frequency change was observed at 29°C, which may suggest that the two stressors, temperature and laboratory environment, act synergistically. Only a small fraction (1%, Fig 2, yellow) of windows did not change in frequency beyond what is expected by drift in either treatment. Qualitatively similar results were obtained when the comparison between the two temperature regimes was performed for single SNPs or averaged across windows of 50, 250 and 500 SNPs as well as with different False Discovery Rate (FDR) thresholds (Fig SI 3).

The elevated selection response to a high temperature laboratory environment may indicate that temperature stress increases the phenotypic variance on which selection can operate. We scrutinized this hypothesis by re-analyzing RNA-Seq data from the two parental genotypes exposed to 18° and 29°C (Chen et al., 2015). Although the difference in gene expression between the two genotypes was much more pronounced at 29°C than at 18°C, we found a positive correlation of the differences in gene expression between the two genotypes between 18°C and 29°C (Fig 2C, Spearman ρ = 0.68). This confirmed that the hot temperature environment amplifies phenotypic differences between the two genotypes. Since traits with a higher phenotypic variance are responding more strongly to selection, we compared the genomic response at the two temperatures and found that the correlated expression changes are mirrored by the parallel selection response of genomic windows at the two temperatures (Fig 2A,2B,2D, Spearman ρ = 0.49). Similar to the parental gene expression with a 36% increase of the median absolute logFC S/O at 29°C, the median absolute AFC increased by 48% at 29°C relative to 18°C.

Given that both gene expression differences and selective responses are correlated between temperatures, we were interested whether they are actually functionally linked. We asked if the genes with the largest temperature-specific expression differences between the two parental genotypes also display the largest temperature-specific selection response. We measured the correlation of the parental gene expression and AFC differentials between 29°C and 18°C (|(Sam-Or)_18°C_ - (Sam-Or)_29°C_|). Neither for the full set of genes nor the top genes (ranked by decreased differential of logFC S/O or AFC), the AFC differential was significantly correlated with the parental expression differential (Fig SI 4). We conclude that the allele frequency changes in the experimental evolution are not primarily driven by parental expression differences. Thus, either parental gene expression differences have limited implication for fitness or the observed gene expression differences are driven by trans-acting factors rather than by cis-regulatory variation. We studied the potential of transcription factors driving the parental expression differences and found that the 50 genes with the strongest expression differences between the parents (ranked by decreasing absolute logFC) at 18°C and at 29°C (17 genes in common) were enriched for many rather than a few transcription factor binding sites (152 at 18°C and 133 at 29°C). We conclude that the temperature-specific gene expression differences between Samarkand and Oregon-R could be driven by many transcription factors, consistent with gene expression having a polygenic architecture.

## DISCUSSION

We studied the selective impact of two different environments on a genomic scale by combining laboratory and temperature adaptation. Contrary to the recommended design for E&R studies (Kofler & Schlötterer, 2014), which facilitate the identification of a moderate number of selection targets occurring at sufficiently high starting frequencies, we did not use a large number of founder genotypes. Rather, we restricted the variation to only two different founder genotypes, as also done in experimental evolution with yeast (e.g. Kosheleva & Desai, 2018). The advantage of this experimental design is that all selection targets have the same starting frequency and a more parallel selection response is expected because polygenic traits have fewer selection targets contributing to a new trait optimum (Barghi & Schlötterer, 2020, Höllinger et al., 2019).

We found pronounced selection responses, which fall into two classes – temperature-specific (change in the direction of the allele frequency change) and laboratory adaptation (parallel selection with similar intensities in the two temperature regimes). In addition, 28% of the genomic windows responded in the same direction, but to a different extent, indicating the joint contribution of both environmental factors.

Temperature-specific adaptation implies that temperature uncovers fitness differences between genotypes. 14% of the genomic windows responded only at 29°C and 8% were private to 18°C, a pattern consistent with conditional neutrality (Schnee & Thompson, 1984). The selection responses private to 18°C indicate that even at an assumed benign temperature, selection occurs - highlighting the challenge of performing control experiments for temperature adaptation. In 10% of the windows, different alleles were favored at each temperature regime. Such temperature-specific selection responses provide an excellent starting point for the identification of causative variants driving temperature adaptation. Nevertheless, the broad genomic regions responding to selection preclude the distinction between causative variants and neutral hitchhikers (Franssen et al., 2015, Nuzhdin & Turner, 2013, Tobler et al., 2014) after 20 generations. Additional generations as well as a larger population size could facilitate the uncoupling of the causative variants from the passenger alleles and improve resolution (Langmüller et al., 2021, Phillips et al., 2020).

Laboratory adaptation is an umbrella term for stressors that can be attributed to the experimental laboratory setup (Matos et al., 2002, Matos et al., 2000, Simoes et al., 2007). Examples of such factors are adaptation to high larval density / early fertility (Hoffmann et al., 2001, Mueller, 1997), sexual selection (Fricke & Arnqvist, 2007) and adaptation to the laboratory food (Bochdanovits & de Jong, 2003, Lai & Schlötterer, 2021, Vijendravarma et al., 2012). With about one third of the genomic windows showing a parallel selection response at both temperature regimes, laboratory adaptation was an important factor in this study.

Of particular interest is the significant difference in allele frequency change for 28% of the windows with parallel selection signatures, because it suggests an interaction between laboratory adaptation which drives the parallel response and temperature which modulates the strength of selection. Adaptation to larval density may be an excellent candidate driving this laboratory adaptation because we maintained the populations at high, but not well-controlled, larval densities. Higher larval density does not only increase competition (Mueller, 1988) but also interactions between larval density and heat stress survival (Arias et al., 2012) as well as body size (James & Partridge, 1998) and locomotor activity (Schou et al., 2013) were previously detected.

Because laboratory experiments cannot fully match natural conditions, it is not possible to conduct these experiments in a full factorial design - we can only modulate the temperature under laboratory conditions, but not in the natural environment. This implies that our design cannot distinguish between additive and interaction effects of temperature *per se* and laboratory adaptation. Selection responses driven by multiple selection factors can be problematic for the interpretation of the selection signatures. Experiments contrasting ancestral and evolved populations cannot distinguish between laboratory adaptation and selection driven by the focal factor (temperature in our study). When populations are compared, which evolved towards two different focal environments (here, different temperatures), the influence of laboratory adaptation is less severe: selection responses with the same direction and magnitude will not be seen in this contrast. Parallel selection responses that differ in magnitude will be interpreted as a pure temperature effect. An experimental design, which does not only include populations evolved in two different focal conditions (*i.e*. different temperatures), but also the ancestral founder populations, similar to this study, can distinguish between laboratory adaptation, adaptation to focal factor and combined effects. Nevertheless, if laboratory adaptation interacts with temperature (or other focal factors), it is possible that small differences in laboratory environment (e.g. food recipe) may result in a different selection response. We propose that this may contribute to the difficulties to replicate temperature-associated effects.

An alternative explanation for the shared directional selection response at 18°C and 29°C is the presence of genotype-specific deleterious mutations. Since the two parental strains were maintained at small effective population size for many generations, it is conceivable that the influence of deleterious alleles is more pronounced than for genotypes freshly collected from wild. The selection signatures may thus also reflect fitness disadvantage of deleterious combinations of parental alleles that can be detected when the two competing genotypes are maintained at large population size. The observation that temperature stress can both increase and decrease the selection response is consistent with previous studies on deleterious mutations (Agrawal & Whitlock, 2010). While frequently the selection response was found to be positively correlated with stress level (e.g. Shabalina et al., 1997, Chu & Zhang, 2021), also the opposite pattern has been observed (Elena & de Visser, 2003, Kishony & Leibler, 2003). Since we cannot determine how much of the parallel selection response can be attributed to deleterious mutations, it is important to realize that we probably overestimate the influence of laboratory adaptation.

One important limitation of this study is the pronounced linkage disequilibrium in the founder population. During 20 generations, too few recombination events occur to break the association between neighboring windows. This is indicated by autocorrelation of allele frequency up to 8Mb. Thus, even though our analyses are based on windows of 250 neighboring SNPs, neighboring windows are still highly correlated. This implies that neighboring windows may exhibit a selection response due to linkage, rather than due to an independent selection target. Different selection intensities will also determine the size of the genomic region affected, leading to a complex interplay between linkage, direction of selection and selection strength. Therefore, the number of windows showing a given selection response may not be an accurate reflection of the number of selection targets with a given behavior. Nevertheless, the prevailing effects of temperature and laboratory adaptation on fitness should be robust against the effects of linkage.

We conclude that E&R experiments starting with strongly reduced genetic variation can provide a powerful approach to study adaptation, in particular when experiments are performed on an environmental gradient (*i.e*. multiple different temperatures). This setup provides new insights into adaptation, in particular when the E&R experiments are performed for more than only 20 generations, since additional generations provide more opportunity for recombination and the selection targets can be characterized with a higher resolution.

## DATA AND SCRIPTS AVAILABILITY

The sequencing data underlying this article are available in the European Nucleotide Archive (ENA) at https://www.ebi.ac.uk/ena/browser/view/, and can be accessed with PRJEB46805 from Burny et al, 2021 (29°C) and XXX for new data (18°C) specifically generated for this study. All scripts (command lines and data analysis) and final files underlying this article are available in Zenodo at https://dx.doi.org/10.5281/zenodo.5614819. Additional table and figures underlying this article are available in its online supplementary material.

## ACKNOWLEDGEMENTS

We thank Sheng-Kai Hsu and Wei-Yun Lai for sharing code for gene ID conversion and for the transcription factor binding site analysis and Anna Maria Langmüller for fruitful discussions. Illumina sequencing of the samples was performed at the VBCF NGS Unit (www.viennabiocenter.org/facilities). This work was supported by the Austrian Science Funds (FWF; grant numbers P29133, W1225). The authors declare no conflicts of interest.

## Notes

### Competing Interest Statement

The authors have declared no competing interest.

