## Supplementary material for "Genome-wide selection signatures reveal widespread synergistic effects of culture conditions and temperature stress in *Drosophila melanogaster*": SI

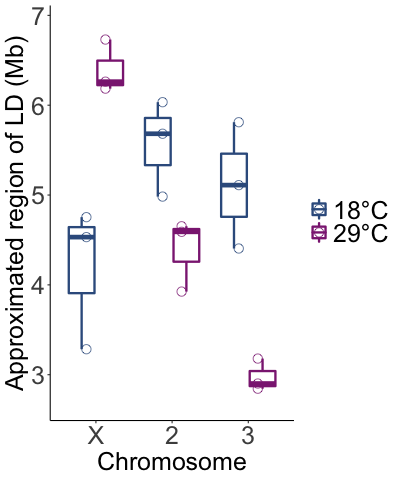

**Figure SI 1.** Large genomic regions respond at both temperatures. We estimated the median size of genomic regions with pronounced linkage at F20 by the autocorrelation in allele frequency between non-overlapping windows of 250 SNPs as in Burny et al, 2021. The distance between windows where the autocorrelation is no longer significant is used as measure of association. In the jittered boxplots, one dot represents one chromosome and a given replicate (18°C, blue; 29°C, purple).

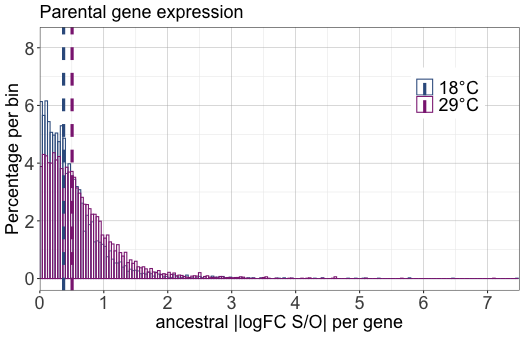

**Figure SI 2.** Histograms of parental gene expression differences per gene at 18°C (blue) and at 29°C (purple). The dashed lines represent at 18°C and 29°C the median absolute logFC S/O (0.37 at 18°C and 0.50 at 29°C, B; p-value paired Wilcoxon one-sided test=5×10^-148^).

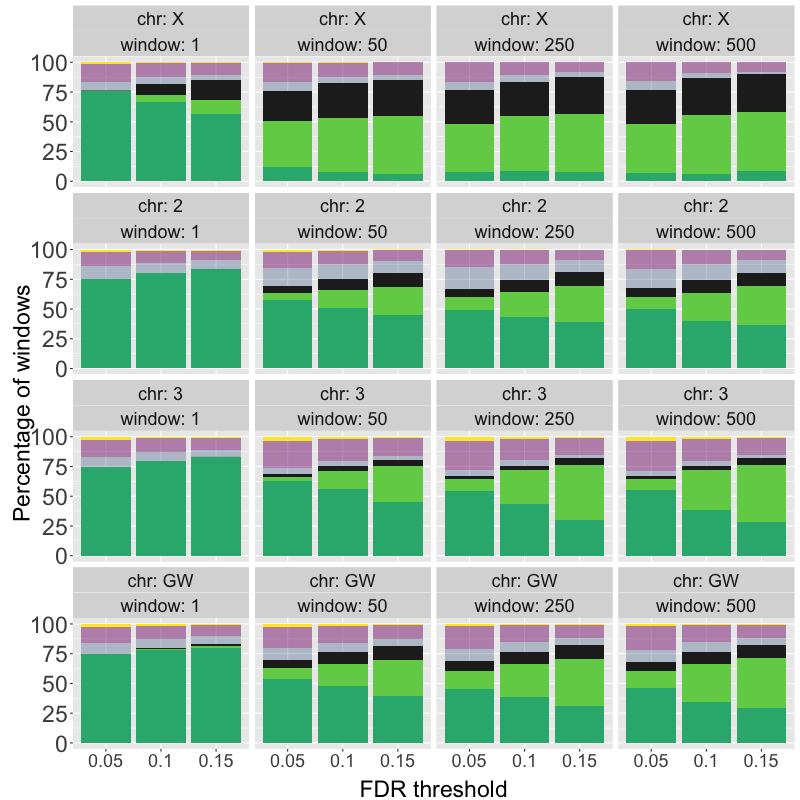

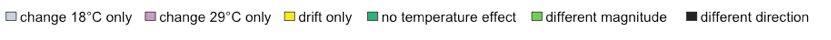

**Figure SI 3**. Influence of the window size (1, 50, 250, 500 SNP(s)) and FDR threshold (x-axis, 0.05, 0.1, 0.15) for the partitioning of the genomic windows´ diagnostic (y-axis, color code indicated in the legend). Individual chromosomes and genome-wide estimates are shown.

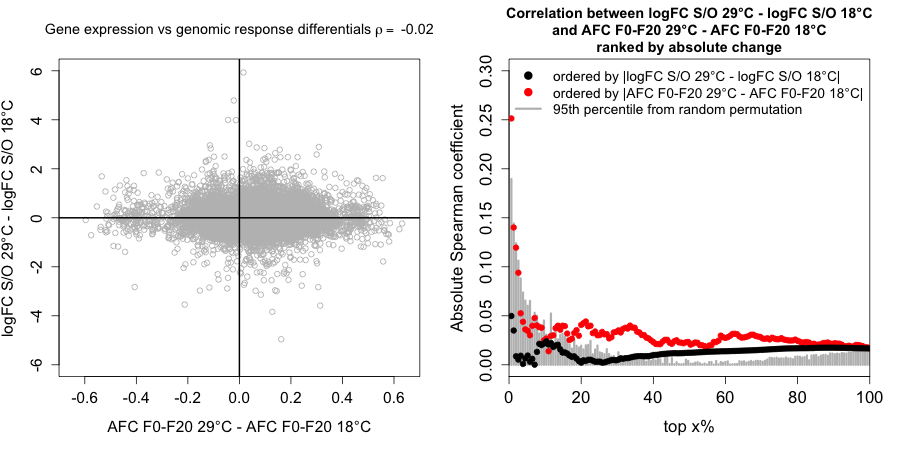

**Figure SI 4**. Correlation between ancestral gene expression and genomic response after 20 generations. *Left.* The temperature effect on the gene expression differences between the two parental genotypes (Samarkand and Oregon-R, y-axis) is plotted against the temperature effect of the allele frequency changes (x-axis). Each data point corresponds to a gene. The Spearman correlation (ρ) coefficient is reported. *Right.* The correlation between gene expression differential (logFC S/O 29°C - logFC S/O 18°C) and allele frequency differential (AFC 29°C - AFC 18°C) are measured in bins containing an increasing number of genes, which are ranked by expression (red) or allele frequency (black) differential.

**Table SI 1.** Partition of the genome per class. The class definition is indicated in column 3 where the conditioning is made on p-values corrected with the Benjamini-Hochberg procedure for either neutrality tests at 18°C and 29°C (p.adj_w_^18°C neutral^ and p.adj_w_^29°C neutral^) or from the linear model on the non-neutral windows (p.adj_w_^LM^) (see Methods). The percentage of windows affected in each class is reported in column 4 per chromosome and average genome-wide (GW), obtained with an False Discovery Rate (FDR) threshold of 5%, 10% and 15% (1^st^, 2^nd^ and 3^rd^ sub-row) and for non-overlapping windows of 250 SNPs.

| Class label | Color code | Logical condition per window w | Percentage of windows ^2^ | | | | |
| --- | --- | --- | --- | --- | --- | --- | --- |
|  |  |  | FDR (%) | 2 | 3 | X | GW |
| Drift only | yellow | p.adj_w_^29°C neutral^ ≥ FDR and  p.adj_w_^18°C neutral^ ≥ FDR | 5  10  15 | <1  <1  0 | 4  2  1 | 0  0  0 | 2  1  <1 |
| Change at 18°C only | light blue | p.adj_w_^29°C neutral^ ≥ FDR and  p.adj_w_^18°C neutral^ < FDR | 5  10  15 | 18  14  10 | 5  5  3 | 7  5  4 | 11  8  6 |
| Change at 29°C only | light purple | p.adj_w_^29°C neutral^ < FDR and  p.adj_w_^18°C neutral^ ≥ FDR | 5  10  15 | 14  12  9 | 24  18  14 | 17  11  8 | 19  14  11 |
| No temperature effect | dark green | p.adj_w_^29°C neutral^ < FDR and  p.adj_w_^18°C neutral^ < FDR and  p.adj_w_^LM^ ≥ FDR | 5  10  15 | 49  43  39 | 54  43  30 | 8  8  7 | 46  38  31 |
| Different magnitude | light green | p.adj_w_^29°C neutral^ < FDR and  p.adj_w_^18°C neutral^ < FDR and  p.adj_w_^LM^ < FDR and  *α_w_^intercept^* and *α_w_^temperature^* of same sign^1^ | 5  10  15 | 11  21  30 | 10  29  47 | 40  46  49 | 14  28  40 |
| Different direction | black | p.adj_w_^29°C neutral^ < FDR and  p.adj_w_^18°C neutral^ < FDR and  p.adj_w_^LM^ < FDR and  *α_w_^intercept^* and *α_w_^temperature^* of different sign^1^ | 5  10  15 | 7  10  11 | 3  4  5 | 28  29  31 | 8  10  11 |

^1^ Due to the linear model formulation, *α_w_^intercept^* and *α_w_^temperature^* of same (different) sign is similar as AFC_w_^18°C^ and AFC_w_^29°C^ of same (different) sign (see Methods).

^2^ Percentages have been rounded.
